# Chemical genomic guided engineering of gamma-valerolactone tolerant yeast

**DOI:** 10.1101/213991

**Authors:** Scott Bottoms, Quinn Dickinson, Mick McGee, Li Hinchman, Alan Higbee, Alex Hebert, Jose Serate, Dan Xie, Yaoping Zhang, Joshua J Coon, Chad L Myers, Robert Landick, Jeff S Piotrowski

## Abstract

**Background:** Gamma valerolactone (GVL) is a promising technology for degradation of biomass for biofuel production; however, GVL has adverse toxicity effects on fermentative microbes. Using a combination of chemical genomics and chemical proteomics we sought to understand the mechanism toxicity and resistance to GVL with the goal of engineering a GVL-tolerant, xylose-fermenting yeast.

**Results:** Chemical genomic profiling of GVL predicted that this chemical affects membranes and membrane-bound processes. We show that GVL causes rapid, dose-dependent cell permeability, and is synergistic with ethanol. Chemical genomic profiling of GVL revealed that deletion of the functionally related enzymes Pad1p and Fdc1p, which act together to decarboxylate phenolic acids to vinyl derivatives, increases yeast tolerance to GVL. Further, overexpression of Pad1p sensitizes cells to GVL toxicity. To improve GVL tolerance, we deleted *PAD1* and *FDC1* in a xylose-fermenting yeast strain. The modified strain exhibited increased anaerobic growth, sugar utilization, and ethanol production in synthetic hydrolysate with 1.5% GVL, and under other conditions. Chemical proteomic profiling of the engineered strain revealed that enzymes involved in ergosterol biosynthesis were more abundant in the presence of GVL compared to the background strain. The engineered GVL strain contained greater amounts of ergosterol than the background strain.

**Conclusions:** We found that GVL exerts toxicity to yeast by compromising cellular membranes, and that this toxicity is synergistic with ethanol. Deletion of *PAD1* and *FDC1* conferred GVL resistance to a xylose-fermenting yeast strain by increasing ergosterol content in cells. The GVL-tolerant strain fermented sugars in the presence of GVL levels that were inhibitory to the unmodified strain. This strain represents a xylose fermenting yeast specifically tailored to GVL produced hydrolysates

## Background

Lignocellulosic biomass derived biofuels and commodity chemicals provide a myriad of sustainable bioproducts. Before biomass can be converted to biofuels, it must be first pre-treated and hydrolyzed to fermentable sugars. However, pre-treatment and hydrolysis processes release fermentation inhibitors, which throttle fermentation rates at a substantial economic cost [1,2].

Fermentation inhibitors come in many forms, and the landscape of these inhibitors is constantly changing as new pre-treatment, hydrolysis, and feedstocks technologies are developed [2]. Enzymatic hydrolysis of biomass releases small acids, phenolics, and furans that are a ubiquitous challenge to bioconversion [1,3]. Chemical hydrolysis methods such as γ-valerolactone (GVL) and ionic liquids offer an enzyme free route to fermentable sugars, but come with their own challenges [4,5]. In addition to the biomass-derived inhibitors, the chemicals used for hydrolysis persist in residual amounts in the hydrolysate, and these compounds are known to be harmful to fermentative microorganisms [6,7]. Further, as these chemical catalysts are used in relatively large amounts during hydrolysis, the residual concentrations are often much higher than the small acidic and phenolic inhibitors generated from the biomass.

GVL is the solvent for a promising, new chemical hydrolysis technology to breakdown the cellulose polysaccharides to fermentable sugar monomers [4]. Advantages of GVL include it is recoverable and renewable, as it is a product of biomass conversion. One challenge of this method is the toxicity of residual GVL to fermentative microbes. GVL is mildly toxic to yeast, but this toxicity may be magnified in combination with other inhibitors and the ethanol produced. As such, engineering GVL-tolerant microbes is a means of overcoming toxicity, minimizing the costs of reagent recovery, and improving biofuels produced via GVL hydrolysis.

Chemical genomics-guided bioengineering has recently been used to develop yeast strains that are highly tolerant of ionic liquids [8]. Herein, we used chemical genomics to discover the genome-wide response to toxicity. Using this information, we identified specific genes that mediate toxicity, and then engineered these specific mutations into an industrially viable, xylose-fermenting strain of *Saccharomyces cerevisiae*. This approach offers a rapid method of tailoring existing strains to specific chemical stressors found in industrial bioconversion.

## Methods

### Chemicals, strains, growth conditions, toxicity screening and IC50 determination

Compounds tested were purchased from Sigma, USA. Cells of *S. cerevisiae* (MATα *pdr1Δ*::*natMX pdr3Δ*::*KI.URA3 snq2Δ*::*KI.LEU2 can1Δ*::*STE2pr-Sp_his5 lyp1Δ his3Δ*1 *leu2Δ*0 *ura3Δ*0 *met15Δ*0), referred to as control strain, were grown in 200 µl cultures at 30 °C in YPD, with a drug or DMSO control. Plates were read on a TECAN M1000 over a 48 h growth period. The specific growth rate was calculated using GCAT analysis software (https://gcat3-pub.glbrc.org/) [9]. When presented, IC_50_ values for growth inhibition were calculated from triplicate 8-point dose curves and SigmaPlot 12.0. When presented, error bars are Mean ± Standard error of the mean of at least 3 replicates.

### Chemical genomic analysis

Chemical genomic analysis of GVL was performed as described previously [8,10]. The tested yeast deletion collection had ~4000 strains using the genetic background described in Piotrowski et al (2017) [11]. We grew cells with 2.3% GVL to mimic the amounts found in GVL produced hydrolysates using 200 µL cultures of the pooled, deletion collection of *S. cerevisiae* deletion mutants was grown in yeast extract (10 g/L), peptone (20 g/L), and 2% galactose (YPGal) medium with 2.3% GVL or a 1% DMSO control in triplicate for 48 h at 30 °C. Genomic DNA was extracted using the Epicentre MasterPure™ Yeast DNA purification kit. Mutant-specific molecular barcodes were amplified with specially designed multiplex primers [20]. The barcodes were sequenced using an Illumina HiSeq. Three replicates of each condition (GVL vs DMSO) were sequenced. The barcode counts for each yeast deletion mutant in the presence of GVL were normalized against the DMSO control conditions to define sensitivity or resistance of individual strains using BEANcounter [12]. To determine a p-value for each top sensitive and resistant mutant, we used the EdgeR package [13,14]. A Bonferroni-corrected hypergeometric distribution test was used to search for significant enrichment of GO terms among the top sensitive and resistant deletion mutants [15].

### MoBY-ORF profiling

MoBY-ORF profiling of GVL was conducted by first generating a pooled collection of the yeast GLBRC-Y133 (henceforth Y133) containing the plasmid collection[16,17]. The plasmid pool for transformation was generated as described previously [16]. For yeast transformation, the plasmids were extracted from 150 mL of *E. coli* culture MAXI Prep. Plasmid was used to transform Y133 via high efficiency LiAc transformation. Transformed yeast were plated to YPD+G418 agar plates and incubated until colonies appeared. A total of 50,000 colonies were washed from the plates using 1X PBS, mixed 1:1 with 50% glycerol, and stored at −80 ºC until use. For MoBY-ORF profiling, 25 mL of media containing YPD+2.5% GVL+ G418 was allowed to degas overnight in an anaerobic chamber, and then inoculated with 100 µL of the transformed yeast pool (n=3). Cells were grown for 48h. Genomic DNA was extracted from 1 mL from each culture using modified mini-prep with zymolyase and glass beads. Gene specific barcodes were amplified, processed, sequenced, and analyzed as described above.

### Growth and sugar conversion experiments

We compared the growth of the Y133 *pad1∆ fdc1∆* to the background strain (Y133) in multiple experimental conditions in flasks: YPXD medium (yeast extract 10 g/L, peptone 20 g/L, glucose 20 g/L, and xylose 20 g/L), with 2.5% GVL (aerobic) or 2% GVL (anaerobic, Coy Chamber); anaerobic synthetic hydrolysate (SynH), containing 60 g/L glucose and 30 g/L xylose)+ 1.5% GVL [3]; aerobic SynH (90g/L glucose and 60 g/L xylose) + 1% GVL. For all experiments, cultures were grown in 25 mL flasks (n=3). Flasks were inoculated with rinsed Y133 or Y133 *pad1∆ fdc1∆* cells to bring the initial OD to approximately 0.1. The flasks were grown for 72 hours with agitation anaerobically at 30 °C. 1 mL samples were taken every 24 hours. Initial and daily samples were measured for OD and submitted for HPLC analysis to quantify sugar consumption and ethanol production [18].

We additionally performed a large scale anaerobic fermentation using 0.5-L bioreactors (BIOSTAT Qplus system from Sartorius, Bohemia, NY, USA) as described in previous work [18]. For these experiments, we used SynH supplemented with FeSO_4_ (20 µM), sodium citrate (10 uM), and GVL (1.5%). Fermentation was done anaerobically by N_2_ sparging and pH was controlled at 5. HPLC samples were collected and analyzed as previously described [18].

### Cell leakage assays

A FungaLight^TM^ Cell Viability assay (Invitrogen L34952) was used to determine if GVL caused membrane damage we used using a Guava Flow Cytometer (Millipore, USA) as described previously [19]. Briefly, log-phase cultures were incubated with 0, 2.5, 5, or 10% GVL or EtOH for 4 h at 30 °C. The cells were then stained and immediately read by flow cytometry to determine the population of stained cells (permeable) vs non-stained cells.

### Synergy screening

To test for synergy, a 6×6 dose matrix was initially used to identify potentially synergistic dose combinations, these points were then confirmed in triplicate. 200 µl cultures were grown with combinations with GVL (2%), ethanol (4%), and the relevant single agent and solvent controls with their OD measured after 24 h. Synergy was determined by comparing actual optical density in the presence of compound combinations to an expected value calculated using the multiplicative hypothesis. This assumes that, in the absence of an interaction, each compound would decrease the OD of the cell culture by the same fraction in the presence of the other compound as it does when applied alone, *i.e.*, *E = A*B/C*, where *E* is the expected OD, *A* is OD when compound A is applied alone, *B* is OD when compound B is applied alone, and C is OD of the control culture (DMSO). In the presence of synergy, the actual OD value is lower than the expected OD. A paired t-test was used to confirm statistical significance of this difference in three replicates of the experiment.

### Determination of inhibitors present in GVL hydrolysates by RP-HPLC-HR/AM-MMS/MS in GVL hydrolysates

GVL hydrolysate samples were diluted 1:10 and 20 µL samples were analyzed by reverse phase (C18) HPLC-high resolution / accurate tandem mass spectrometry (HPLC-MS/MS). Peak areas of peaks matching in retention times and accurate mass +/-10 ppm MS/MS transitions of authentic reference standards were used to calculate concentrations by comparison to an external standard curve as described previously [18].

### Chemical proteomics

For yeast proteomics, triplicate 10 mL of YPD + 1 % GVL or YPD were inoculated with the control strain to a starting OD_600_ of 0.01 and incubated at 30 °C with shaking at 200 rpm. 2 mL of each culture was harvested when they reached an OD_600_ of ~0.5 (mid log phase growth). Cells were pelleted at 10,000 rpm, the media removed, and stored at −80 °C until processing for proteome analysis.

Yeast cell pellets were resuspended in 6 M GnHCl (Sigma, St. Louis, MO) with 50 mM Tris pH 8.0 (Sigma, St. Louis, MO), boiled for 5 min, and precipitated by adding methanol (Thermo Fisher Scientific, Pittsburgh, PA) to a final concentration of 90 %. The precipitate was centrifuged at 10,000 rcf for 5 min, decanted, and air dried. The protein pellet was resuspended in 8 M urea (Sigma, St. Louis, MO) with 100 mM Tris pH 8.0, 10 mM Tris (2-carboxyethyl) phosphine (Sigma, St. Louis, MO), and 40 mM chloroacetamide (Sigma, St. Louis, MO). The resuspended sample was diluted to 1.5 M urea with 50 mM Tris pH 8.0. Trypsin was added to a final ratio of 1:20 (enzyme to protein) and the samples were incubated at ambient temperature overnight. Peptides were desalted over Strata-X cartridges (Phenomenex, Torrance, CA). Desalted peptides were dried in a speed vac and resuspended in 0.2 % formic acid (Thermo Fisher Scientific, Rockford, IL). Peptides were quantified with the Pierce quantitative colorimetric peptide assay kit (Thermo Fisher Scientific, Rockford, IL).

For each analysis, 2 µg of peptides were separated across a 30 cm, 75 µm i.d. column packed with 1.7 µm BEH C18 particles (Waters, Milford, MA). Mobile phase A was 0.2 % formic acid and B was 0.2 % formic acid, 70 % ACN, and 5 % DMSO (Thermo Fisher Scientific, Pittsburgh, PA). The gradient was 5–50 % B over 100 min followed by a 100 % B wash and re-equilibration with 0 % B. Eluted peptides were analyzed on a Thermo Fusion Orbitrap (Thermo Fisher Scientific, San Jose, CA). Orbitrap survey scans were performed at 60,000 resolution, followed by ion-trap ms/ms analyses of the most intense precursors (with z = 2–6) for less than 3 s and using a dynamic exclusion of 15 s. The maximum injection time for each ms/ms was 25 ms and the ion-trap resolution was set to turbo.

Peptides were identified and quantified from the MS data using the MaxQuant software suite with the Andromeda and MaxLFQ search and quantitation algorithms, respectively. Spectra were searched against a Uniprot human proteome and common contaminant database concatenated with the reverse sequences. Match between runs was toggled on with the default settings. Peptide and protein identifications were filtered to 1 % FDR, and proteins were quantified by the MaxLFQ algorithm using the default settings. Data was visualized in Spotfire 5.5.0 (TIBCO, USA). A Bonferroni-corrected hypergeometric distribution test was used to search for significant enrichment of GO terms among the top 15 sensitive/resistant deletion mutants with a p value of p < 0.01 [15].

### Ergosterol quantification

The strains used in this study were Y133 and Y133 *pad1Δfdc1Δ*. Yeasts were cultured in triplicate 10 mL of synthetic hydrolysate (SynH) media with and without 1% GVL and incubated on a rotary shaker at 30 °C at 225 rpm until an OD600 of 1.0 was reached. The entire culture was sampled without separation of the cells from the culture medium. The results were obtained by a simple method including saponification with methanolic potassium hydroxide, extraction of non-saponified lipids, derivatization and the ergosterol content was compared to an analytical reference standard.

Yeast cultures were diluted to O.D. 0.1-0.3 with SynH. Cultures were saponified by adding methanolic potassium hydroxide to bacterial culture containing cholesterol and dueterated 5alpha Cholestan as ISTD’s [20].

Variations of this procedure have been used for many years to extract most lipid classes into essentially nonpolar chloroform solvent [21,22]. The extraction solvent (methanol:chloroform) was added to the hydrolyzed culture sample and the original sample tube was rinsed with water and transferred to a separation funnel. Two phases were observed and the lower phase (chloroform) containing the lipids was collected. The original sample tube was rinsed with chloroform and transferred to the separation funnel and the phases were allowed to separate. The lower phase was added to the initial collected chloroform phase and the upper phase was discarded. The total sample volume collected was dried in a heat block under a gentle stream of nitrogen.

Chloroform was added to all dried sample and standard tubes, mixed and aliquoted for derivatization. MSTFA and anhydrous pyridine were added to all samples and standards, heated, cooled, and transferred to an autosampler vial and were immediately analyzed.

Ergosterol standards with a calibration range of 0.194 µg/mL to 396.65 µg/mL were derivatized with MSTFA and anhydrous pyridine. Cholesterol and dueterated 5alpha Cholestan were used as internal standards, and added to all standard levels.

Samples were analyzed on an Agilent technologies 5975C inert XL MSD with Triple-Axis Detector and an Agilent Technologies 7890A gas chromatograph coupled with a CTC Analytics COMBIPAL Autosampler.

The primary column was HP-5MS 5% Phenyl Methyl Silox: 30m × 250um × 0.25um. Carrier Gas was He, 1.2 mL/min with an Inlet temperature of 150 ºC. Primary GC oven program: initial temp of 1500 ºC, hold for 1min; increase 40 ºC per minute to 280 ºC and hold for 0 minutes then 5 ºC per minute to 320 ºC for 1 minute, Deactivated glass split universal liner with glass wool, split ratio 10:1, Quad temp 150 ºC, Source temp 230 ºC. Integration of peak areas, calculation of standard curves and interpolation of concentrations were performed with vendor supplied software; MassHunter Workstation Software B.06.00

## Results

### GVL is the primary inhibitor found in GVL hydrolysates

GVL-produced hydrolysates remain largely unstudied; thus, our first goal was to identify the primary inhibitors of GVL hydrolyates. LC/MS of hydrolysates revealed that three inhibitory compounds were highly abundant in the GVL hydrolysates: GVL, levulinic acid, and hydroxymethylfurfural (HMF); other lignocellulosic derived inhibitors were present, but at much lower concentrations (**Supplementary Information 1**). Currently, GVL hydrolysates contain a high level of residual GVL (~250 mM); at this level, GVL is the most abundant inhibitor in GVL hydrolysates with half-maximal inhibitory concentration (IC_50_) of 270 mM (**Fig. 1**). Based on this result, we focused on understanding GVL toxicity and developing a tolerant yeast strain.

**Figure 1.**
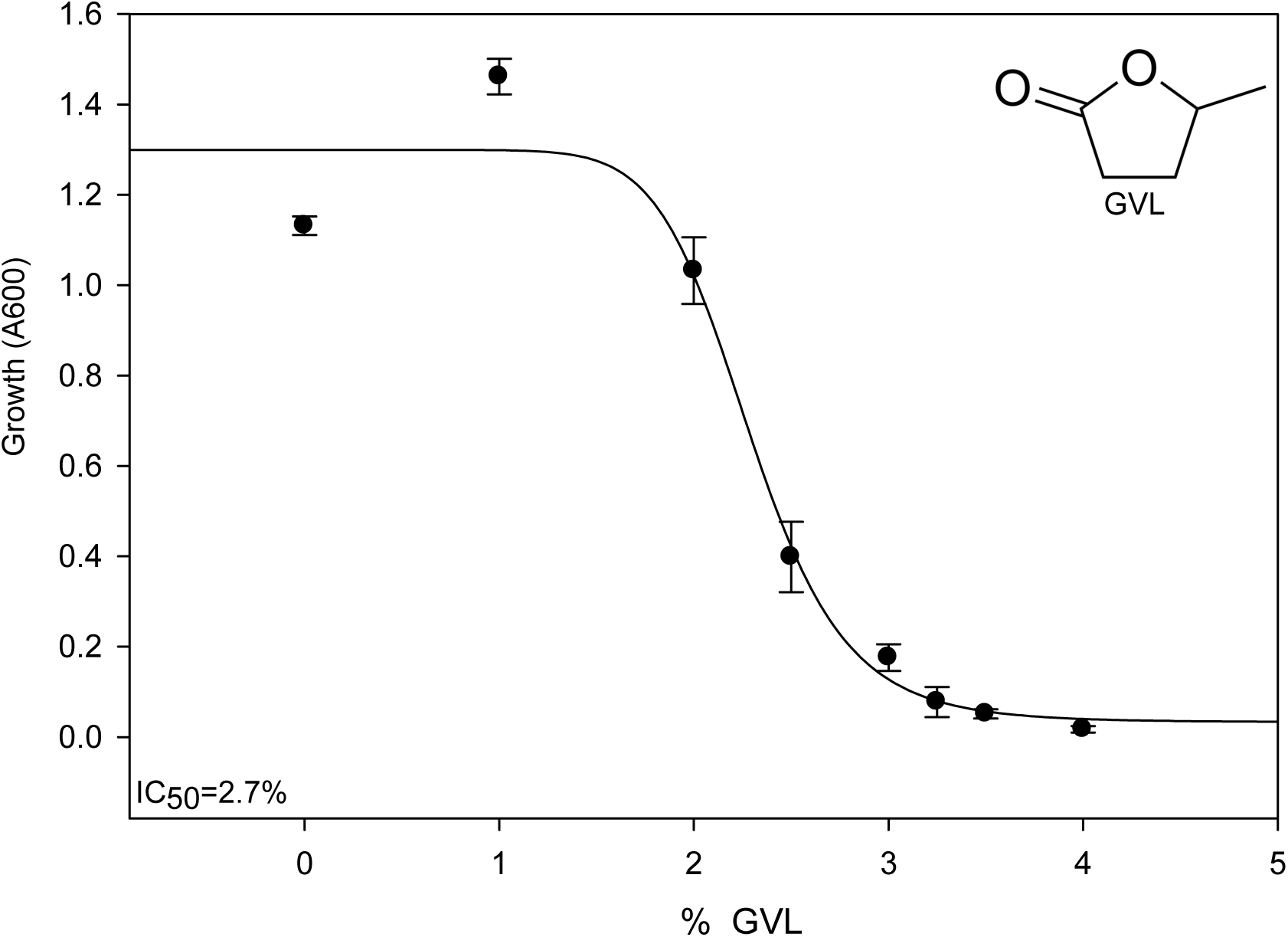
Production of GVL hydrolysates and toxicity. The half-maximal inhibitory concentration (IC50) of GVL against our control strain in rich media was estimated using an 8 pt dose curve (Mean±S.E, n=3).

### Chemical genomic predicts GVL targets cellular membranes and membrane-bound processes

To understand the mode of action of GVL toxicity we conducted chemical genomic analysis. This is a reverse genetics method that uses collections of defined gene mutants, and uses the response of these mutants in the presence of a chemical stress to gain functional insight into the chemical’s mode of action and cellular target [23]. We first tested the yeast deletion collection with standard rich medium with 2% galactose (YP-Gal) containing 2.3% GVL and used barcode sequencing to identify the fitness response of the individual deletion mutants.

We identified 843 significantly responsive deletion mutants (p<0.01, **Supplementary Information 2**). Among the top 10 sensitive deletion mutants, we found significant enrichment for genes involved in late endosome to vacuole transport (p<0.01, **Fig. 2a**), driven by deletion mutants of *SEC28*, *VPS38*, *DID2*. We validated the mutants within this Gene Ontology (GO) category by testing single mutants individually, and found all gave a lower IC_50_ compared to the control strain (**Fig. 2b**). Deletion mutants of these 3 genes exhibit increased sensitivity to ethanol, heat, the ionophore nigercin, and the ergosterol biosynthesis inhibitor miconazole [24–29]. When we correlated the chemical genomic profile of GVL with the yeast genetic interaction network database [30], we found significant enrichment for genes involved in golgi-vesicle mediated transport among the top 10 correlations (p=0.001). *RET2* was consistently predicted as the top correlation for the GVL chemical genomic profile. Ret2p is a subunit of the coatomer complex involved in retrograde transport between Golgi and ER is also involved in golgi transport of vesicles [31]. *RET2* mutants similarity show increased sensitivity to heat and membrane disrupting agents [24,26]. We correlated the chemical genomic profile of GVL to existing chemical genomic datasets [23], and found its profile was significantly similar to profiles of the ionophore nigericin (p<0.01) and the phosphotidylserine targeting agent papuamide (p<0.01), [23]. Taken together, these data suggest GVL could exert toxicity by damaging membrane integrity.

**Figure 2.**
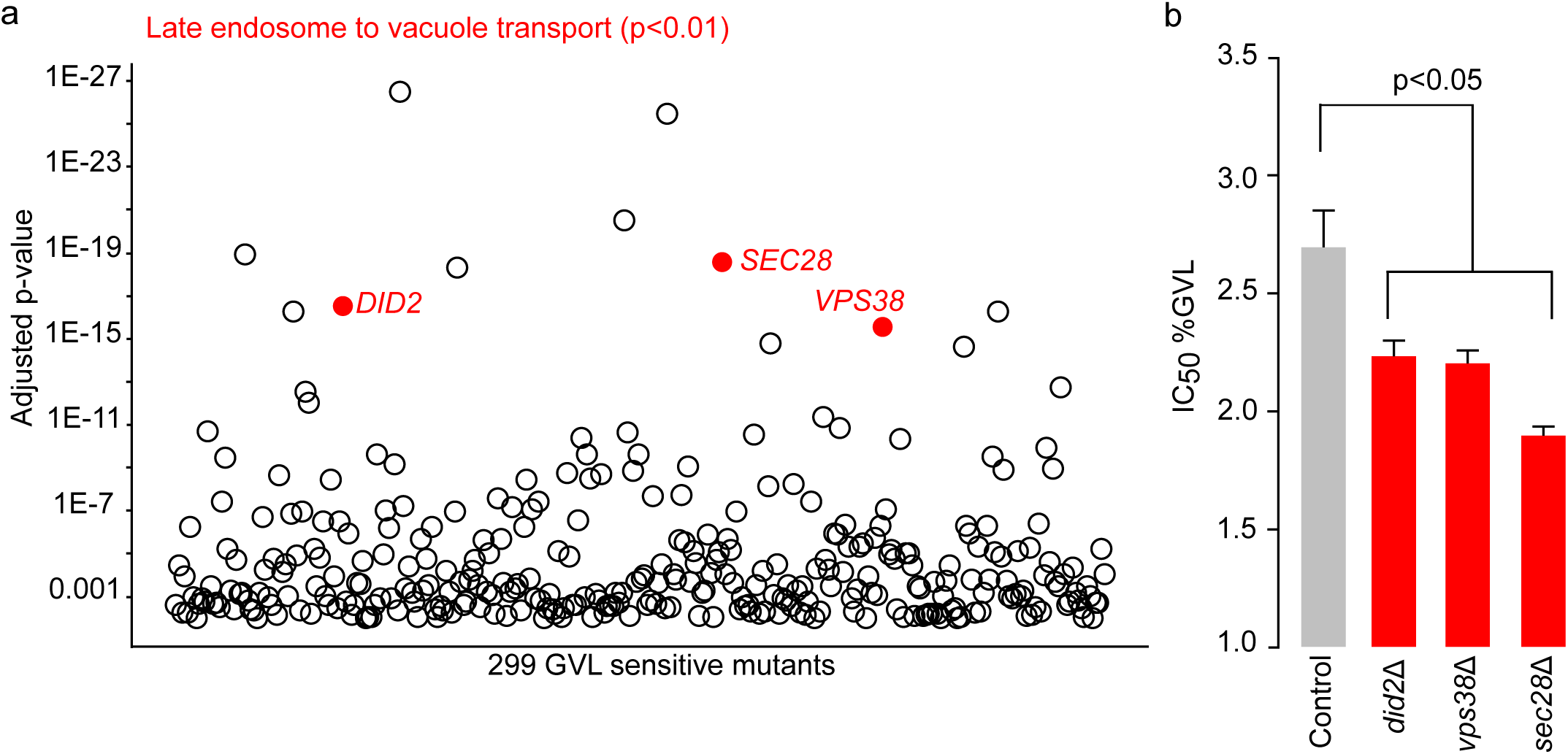
Chemical genomic profiling of GVL. Chemical genomic profiling of the yeast non-essential deletion mutant collection was used to identify mutants significantly sensitive to GVL (**a**, n=3). The p-value of mutants with a decreased fitness in the presence of 2.3% GVL compared to blank medium is plotted on the Y-axis. Of the top 20 significantly responsive mutants, strains annotated to the GO process Late Endosome to Vacuole Transport (GO:0045324) are presented in red. The GVL IC_50_ of *did2Δ, vps38Δ*, and *sec28Δ* versus the control strain was estimated in rich medium (**b**, Mean±S.E, n=3). ANOVA was used to detect differences in growth.

### GVL damages membranes and is synergistic with ethanol

To confirm if GVL treatment can rapidly affect cell integrity, we assessed cell permeability after GVL treatment. Using flow cytometry combined with a dye that is only taken up by cells with damaged membranes, we found a rapid and dose dependent effect of GVL on permeability (**Fig. 3a,b**), similar to the effects of ethanol but with a greater magnitude (**Fig. 3b**). Given that both GVL and ethanol can damage cellular membranes, we also tested if these compounds are synergistic. We found a strong synergism between GVL and ethanol in both our lab strain and xylose fermenting strain (**Fig. 3c**). At a 2% GVL concentration and 4% ethanol concentration, we observed significant synergism in membrane damage between GVL and ethanol (p<0.01). This suggest that as ethanol titers increase during fermentation, the toxic effects of GVL and ethanol will magnify each other, which ultimately will affect yield.

**Figure 3.**
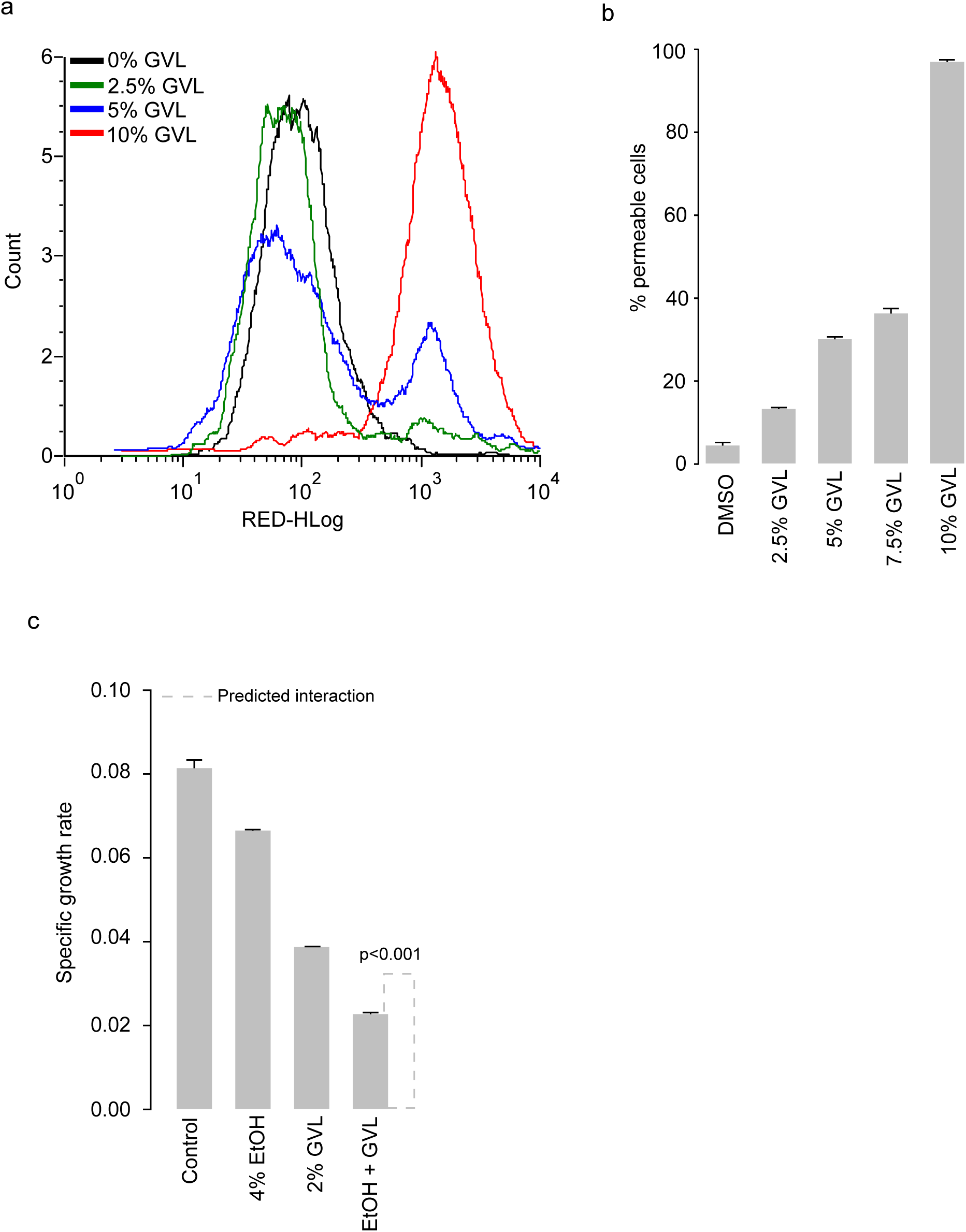
The effects of GVL on cell permeability and synergism with ethanol. Cells were treated with GVL for 4 h GVL, then incubated with a dye that is only taken up by cells with compromised membrane integrity, which results in increased red fluorescence (**a**). The percent of permeable cells in populations of cells treated with either GVL or EtOH (**b**, Mean±S.E, n=3). Specific growth rate of cells treated with 2% GVL, 4% EtOH, or a combination of both (**c**, Mean ± S.E, n=3), The dashed line indicated the predicted level of inhibition if GVL and EtOH were additive in their toxicities.

### Deletion of Pad1p and Fdc1p enhance GVL tolerance

Importantly for our goal, we also investigated gene deletions that increased resistance to GVL. Among the top GVL-resistant mutants we found a significant enrichment for genes involved in phenylpropanoid metabolic process (p<0.002, **Fig. 4a**), driven by deletion mutants of *PAD1* and *FDC1*. Single mutant validations reveal deletion of these genes improved GVL tolerance (**Fig. 4b**). Pad1p is responsible for generating a co-factor needed by Fdc1p to decarboxylate phenolic acids [32–34].

**Figure 4.**
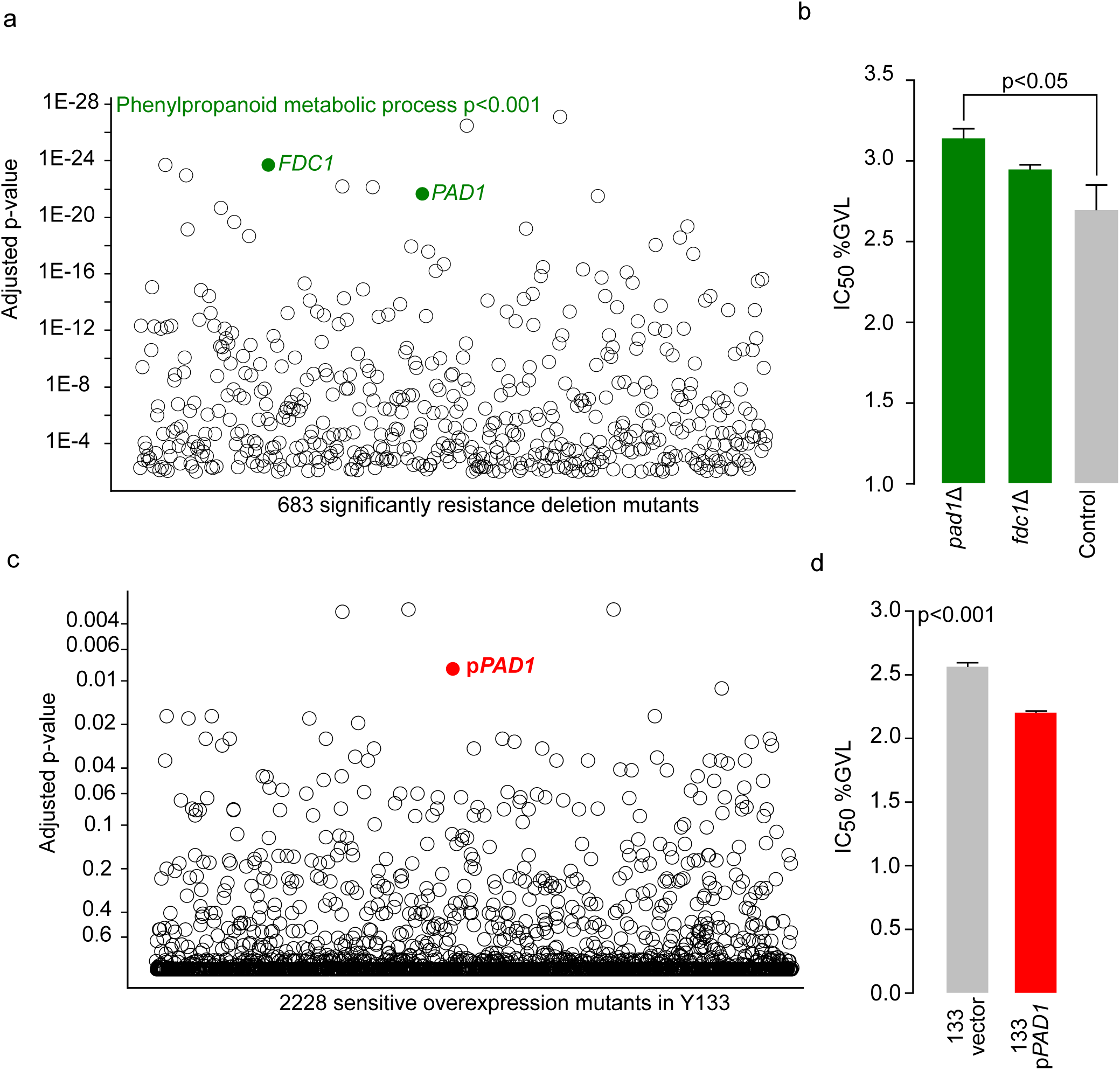
Identifying genes mediating GVL toxicity by deletion and overexpression mutant profiling. The p-value of mutants with an increased fitness in the presence of 2.3% GVL compared to blank medium is plotted on the Y-axis (**a**, n=3). Of the top 20 significantly responsive mutants, strains annotated to the GO process Late Phenylpropanoid metabolic process (GO:0009698) are presented in green. GVL IC50 values of *pad1Δ* and *fdc1Δ* versus the control strain (**b**, Mean ± S.E, n=3). Using the xylose fermenting yeast Y133 transformed with the 2 µ barcoded plasmid collection (MoBY 2.0), we identified genes that confer GVL sensitivity when overexpressed (**c**, n=3). The p-value of overexpression mutants with a decreased fitness in the presence of 2.3% GVL compared to blank medium is plotted on the Y-axis, and the strain harboring p*PAD1* is highlighted in red. GVL IC50 values of p*PAD1* versus an empty vector (**d**, Mean ± S.E, n=3).

### Overexpression chemical genomic profiling confirms Pad1p mediates GVL toxicity

We wanted to extend our chemical genomic analysis to an industrially relevant, xylose ferment yeast strain. At present, there are no available genome-wide deletion mutant collections in industrial yeast, so we took a complementary approach using overexpression plasmids that could be introduced to any yeast. The MoBY-ORF 2.0 plasmid collection, which encodes barcoded versions of 95% of all *S. cerevisiae* genes each expressed on a 2µ plasmid [35], enabled this approach. This collection of plasmids can be pooled and transformed into any yeast to allow investigations of the effect of gene dose under stress conditions. We transformed a version of the xylose-fermenting yeast Y133 [36] *en masse* with the pooled plasmid collection and selected over 50K individual transformants (10X genome coverage). We grew this pooled transformant collection in the presence of 2.5% GVL or a water control under anaerobic conditions in glucose/xylose-containing media and assessed the effects of increased gene dose on growth in the presence of GVL. We found the Pad1p overexpression mutant was one of the top sensitive strains (p<0.01, **Fig. 4c**). We confirmed with single mutant cultures that overexpression of *PAD1* causes GVL sensitivity. The IC_50_ of Y133+p*PAD1* was 2.2 %, compared to 2.56 % of vector control (**Fig. 4d**, p<0.001).

### Deletion of *PAD1* and *FDC1* in a xylose-fermenting strain confers GVL tolerance

Chemical genomic profiling and validation of individual mutants confirmed that the Pad1p and Fdc1p were involved in GVL toxicity. We chose to engineer these deletions into a xylose-fermenting yeast strain Y133. *PAD1* and *FDC1* are adjacent on chromosome IV, and as such we were able to delete both genes at the same time using a single transformation with a PCR product of the antibiotic resistance marker KanMX flanked by homologous regions upstream of *PAD1* and downstream of *FDC1* (**Fig. 5a**). We confirmed deletion of both genes by PCR (**Fig. 5b**).

**Figure 5.**
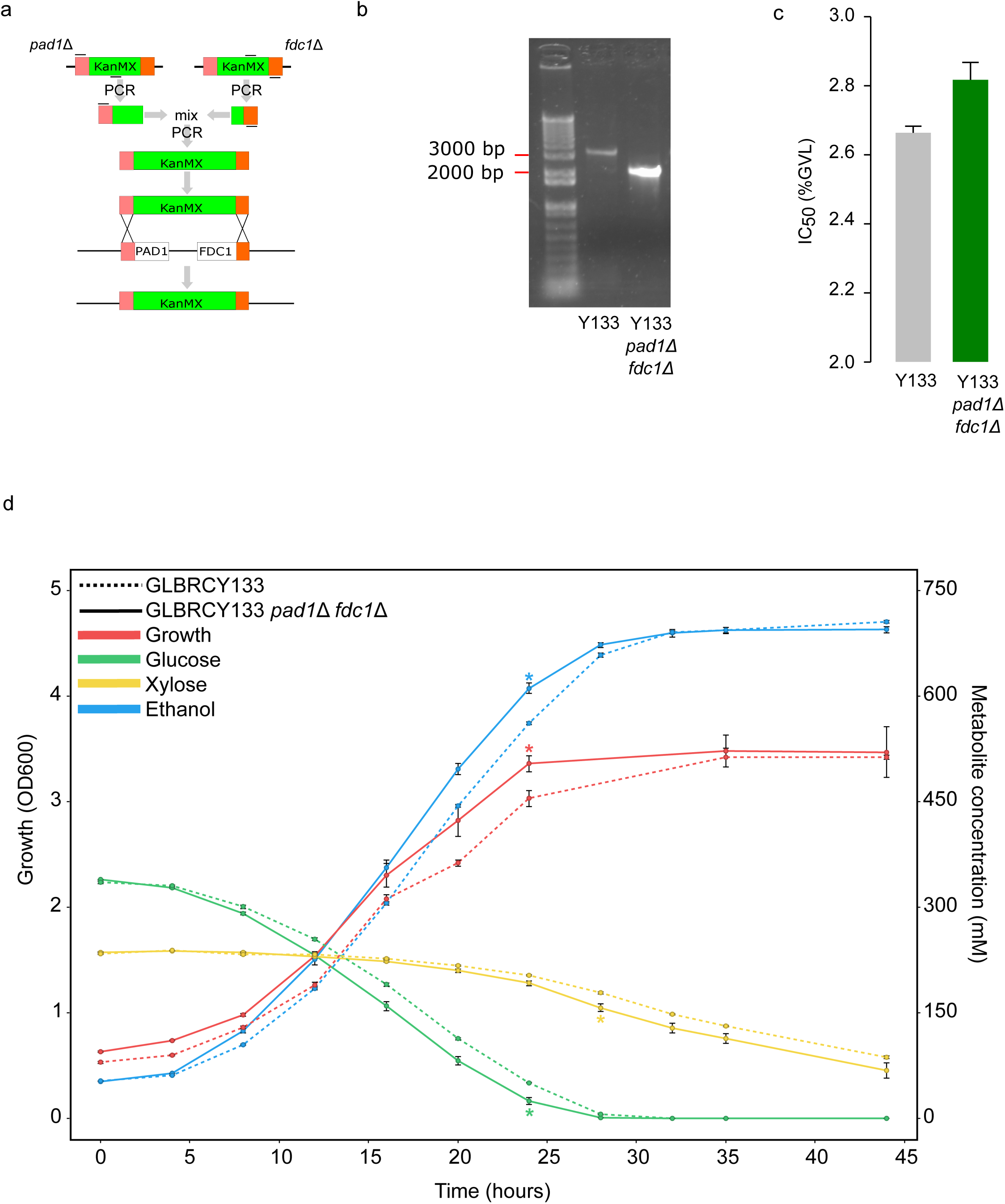
Deletion of *PAD1* and *FDC1* and the effect of this modification on GVL tolerance in a xylose fermenting yeast. We used a two-step PCR approach to simultaneously delete *PAD1* and *FDC1* in Y133, which are adjacent on chromosome IV (**a**), and confirmed deletion by PCR (**b**). GVL IC50 values of Y133 *pad1Δ fdc1Δ* versus Y133 (**c**, Mean ± S.E, n=3). Growth, sugar consumption, and ethanol production of Y133 *pad1Δ fdc1Δ* versus Y133 in synthetic hydrolysate medium containing 1.5% GVL under anaerobic conditions (**d**, Mean ± S.E, n=3) “<” indicates where the engineered strain has significantly greater growth or ethanol production, or significantly less remaining sugars (p<0.05).

The IC_50_ concentration of GVL for the Y133 *pad1∆ fdc1∆* strain was significantly higher than for the Y133 parent (**Fig. 5c**, p<0.01). We tested for GVL tolerance in flask experiments under anaerobic and aerobic conditions in both rich and defined media (**Supplementary Information 3a-d**). In all conditions, the Y133 *pad1∆ fdc1∆* mutant had greater growth than the Y133 strain. Additionally, the tolerant strain had greater initial glucose/xylose use, and ethanol production under anaerobic conditions. Finally, we tested the performance of the Y133 *pad1∆ fdc1∆* strain in a scaled-up fermentation experiment, under industrially relevant anaerobic conditions in a synthetic hydrolysate containing 1.5% GVL. The engineered grew, consumed sugar, and produced significantly more ethanol after 24 h, and had consumed significantly more xylose after 28 h (p<0.05) (**Fig. 5d**).

### Chemical proteomic profiling reveals over production of ergosterol genes in response to GVL in the Y133 *pad1∆ fdc1∆* background

To further understand the mechanism of GVL tolerance of the Y133 *pad1∆ fdc1∆* strain, we used quantitative proteomics to identify how GVL treatment alters protein abundance in the Y133 and Y133 *pad1∆ fdc1∆* strain (**Supplementary Information 4**). Among the top 20 proteins that had increased abundance in the Y133 *pad1∆ fdc1∆* versus the Y133 background (**Fig. 6a**), we observed a significant enrichment for proteins involved in ergosterol (Erg11p, Erg5p, Hmg1p, P=0.009). Interestingly, among the top 20 proteins that had decreased abundance in the presence of GVL (**Fig. 6a**), the Y133 *pad1∆ fdc1∆* strain had significantly enrichment in for genes involved in protein folding (Hsp26p, Hsp82p, Fes1p, Sse2p, Ssa3p, Ssa4p; P=9.2e^−5^), which suggests GVL may further exert toxicity by affecting protein folding in non-resistant strains.

**Figure 6.**
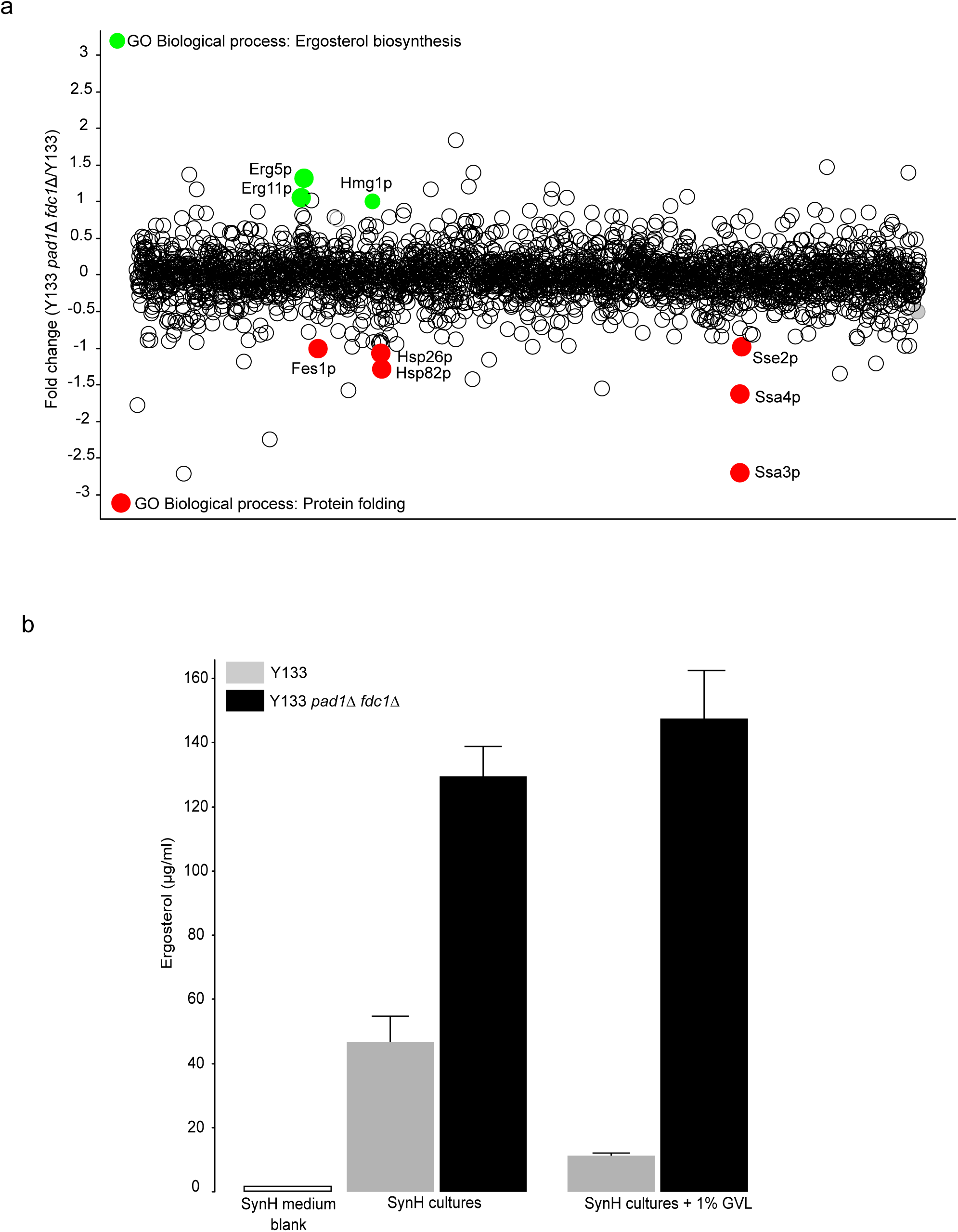
Chemical proteomics and ergosterol quantification for Y133 and Y133 *pad1Δ* and *fdc1Δ*. Protein abundance changes of Y133 *pad1Δ fdc1Δ* versus Y133 in the presence of 1% GVL (**a**, n=3). Fold change values are presented on the Y-axis. Proteins that had increased abundance in the Y133 *pad1Δ fdc1Δ* strain compared to Y133 that are annotated to Ergosterol biosynthesis (GO:0006696) are highlighted in green. Proteins that had decreased abundance in the Y133 *pad1Δ fdc1Δ* strain compared to Y133 that are annotated to Protein folding (GO:0006457) are highlighted in red. Total cellular ergosterol quantification of Y133 *pad1Δ fdc1Δ* and Y133 grown +/-1% GVL (**b**, Mean ± S.E, n=3).

### Y133 *pad1∆ fdc1∆* produces more ergosterol in response to GVL

Ergosterol is a fungal membrane sterol that determines membrane fluidity, and by extension can affect thermal tolerance and tolerance of solvents such as ethanol [37]. Given that proteins involved in ergosterol biosynthesis have increased abundance in the Y133 *pad1∆ fdc1∆* strain compared to the background, we wanted to test in the engineered, GVL-tolerant strain had greater ergosterol content than the parent strain. In untreated SynH medium, the engineered strain produced significantly more ergosterol compared to the parent strain (**Fig. 6b**). This difference was magnified in the presence of 1% GVL, where the engineered strain significantly increased ergosterol production, whereas the parent strain had decreased ergosterol content (**Fig. 6b**).

## Discussion

Using chemical genetics, we discovered that deletion of the functionally related genes *PAD1* and *FDC1* genes confers resistance to GVL in a xylose-fermenting yeast strain, presumably by increasing ergosterol production. Over expression of these genes have been shown to confer resistance to the lignocellulose-derived inhibitors cinnamic acid and ferulic acid [38–40]. Considering this result, yeast harboring *PAD1*/*FDC1* overexpression systems may be incompatible with GVL-produced hydrolysates, and the biocatalyst used should be tailored to the hydrolysate to minimize the effects of process-derived inhibitors (*e.g.*, GVL, ionic liquids), lignocellulose derived inhibitors (*e.g.*, phenolic acids), and end-product inhibitors (*e.g.*, ethanol, isobutanol).

Little is known about the functions of *PAD1* and *FDC1* in yeast, aside from their role in detoxifying aromatic carboxylic acids in lignocellulosic hydrolysates. The bacterial homologs *ubiX* and *ubiD* are known to function in ubiquinone biosynthesis. *PAD1* has a significant negative genetic interaction with *COQ2*, the second step of ubiquinone biosynthesis[41]. Recent work has revealed that *PAD1* is not a decarboxylase as originally described, but rather synthesizes a novel co-factor (a prenylated flavin) required for the decarboxylase activity of *FDC1* [32–34]. *PAD1* and *FDC1* may function in ubiquinone biosynthesis [42], but they are not essential to the process [39].

The relationship between *PAD1*/*FDC1* and ergosterol content also remains unclear. Ubiquinone and ergosterol share the same precursors, as both are products of the mevalonate pathway [43]. Deletion of *PAD1* could reduce flux towards ubiquinone biosynthesis, resulting in greater availability of ergosterol biosynthetic precursor, or induce increased sterol uptake. Additionally, *FDC1* has the most similar genetic interaction profile as *YML082C*, which is regulated by *UPC2*, a gene involved in sterol uptake and biosynthesis [41,44]. Our results are consistent with previous studies that found the laboratory strain CEN.PK113-7D harbored a loss of function mutation, (TAT → TAG, Stop codon) in *PAD1*, and that this strain contained more ergosterol during logarithmic growth [45]. CEN.PK113-7D also harbored mutations in the ergosterol biosynthetic genes *HMG1*, *ERG8*, and *ERG9*, and it is not known if these mutations were secondary to the *PAD1* loss-of-function mutation. Our proteomic analysis revealed the *pad1Δ*/*fdc1Δ* mutant had increased abundance of Hmg1p, Erg5p, and Erg11p in the presence of GVL (**Fig. 6a**). Given that ergosterol biosynthesis is oxygen dependent, and GVL tolerance is apparent in both aerobic and anaerobic media (**Fig. 5d, Supplementary Information 3**), deletion of *PAD1* and *FDC1* may lead to sterol uptake and/or biosynthesis, but sterol uptake would explain tolerance in anaerobic conditions. Further work is needed to more completely characterize the relationship between *PAD1*/*FDC1* and sterol content in yeast.

## Conclusions

We have elucidated the mechanism of -valerolactone toxicity in *Saccharomyces cerevisiae*. Like ethanol, the non-polar solvent GVL compromises membrane integrity, leading to cell lysis. Further, GVL is synergistic with ethanol in toxicity, thus reducing the amount of residual GVL in GVL-produced hydrolysates is critical for not only GVL recovery economics, but also ensuring maximum ethanol titers during fermentation. Given the general effects of GVL on membranes, it is likely that this mechanism of toxicity is similar in prokaryotic biocatalysts.

## List of abbreviations

(GVL): Gamma valerolactone

(YPD): Yeast extract Peptone Dextrose

(SynH): Synthetic hydrolysate

(DMSO): Dimethyl sulfoxide

## Declarations

### Ethics approval and consent to participate

Not applicable

## Consent for publication

Not applicable

## Availability of data and material

Chemical genomic and chemical proteomic data generated or analyzed during this study are included in this published article as Supplementary Information.

## Authors' contributions

QD, SB, LH, AH, JP performed experiments. RL designed experiments and edited the manuscript. CLM developed software for analysis. AH, and JC performed proteomics and analysis. JS, DX and YZ performed large-scale fermentation. AH and MM provided ergosterol quantification and analysis of GVL hydrolysates. JP designed experiments, performed analysis, and wrote the manuscript.

## Authors’ information

JP, QD, SB, LH, AH, JS, DX and YZ are scientists at the University of Wisconsin-Madison and the Great Lakes Bioenergy Research Center (GLBRC). CLM is a Professor in the Department of Computer Science and Engineering at the University of Minnesota. JC is a Professor in the Department of Chemistry at the University of Wisconsin-Madison. RL is a Professor of Biochemistry and Bacteriology at the University of Wisconsin-Madison.

## Acknowledgements

We thank Jim Dumesic and Jeremy Luterbacher for providing samples of GVL hydrolysates and Charlie Boone for providing the yeast deletion collection and the MoBY-ORF 2.0 collections. We thank Trey Sato for critical discussions of this work.

## Additional files

**Additional file 1 –** GVL hydrolysate inhibitors

**Additional file 2 –** Chemical genomic profile of GVL

**Additional file 3 –** Growth, sugar consumption, and ethanol production of Y133 *pad1∆ fdc1∆* vs Y133 grown in anaerobic and aerobic conditions with either rich media (YPXD) or synthetic hydrolysates (n=3, Mean±S.E.).

**Additional file 4 –** Proteomic profile of Y133 *pad1∆ fdc1∆* vs Y133 grown in YPD medium plus 1% GVL

